# RNA genome expansion up to 64 kb in nidoviruses is host constrained and associated with new modes of replicase expression

**DOI:** 10.1101/2024.07.07.602380

**Authors:** Benjamin W. Neuman, Alexandria Smart, Josef Vaas, Ralf Bartenschlager, Stefan Seitz, Alexander E. Gorbalenya, Neva Caliskan, Chris Lauber

## Abstract

Positive-strand RNA viruses of the order *Nidovirales* with genomes larger than ∼20 kb, including the largest known 36.7 kb RNA genome in vertebrate viruses, encode a proofreading exoribonuclease (ExoN). Here, we assemble 76 genome sequences of invertebrate nidoviruses from >500.000 published transcriptome experiments and triple the number of known nidoviruses with >36 kb genomes, including the largest known 64 kb RNA genome. We classify multi-cistronic ExoN-encoding nidoviruses into five groups, according to canonical and non-canonical modes of viral polymerase expression by ribosomes and genome segmentation. The largest group employing the canonical mode comprises invertebrate and vertebrate nidoviruses, including coronaviruses, with genomes ranging from 20-to-36 kb. Four groups with non-canonical expression modes include giant invertebrate nidoviruses with 31-to-64 kb genomes, some of which utilize dual ribosomal frameshifting that we validate experimentally. Thus, expansion of giant RNA virus genomes, the vertebrate/invertebrate host division, and the control of viral replicase expression are interconnected.

## Main

Diverse RNA viruses with no DNA stage in their life cycle have been detected in all eukaryotic and bacterial life forms. Based on genome type, they belong to either positive-strand (ssRNA+), negative-strand (ssRNA-) or double-strand RNA (dsRNA) viruses^1^. Their average genome size is around 10-11 kb and only few lineages have evolved RNA genomes exceeding 20 kb, according to our analysis of the curated RefSeq database^2,3^ and several recent large-scale metagenomics studies that define the current sampling of the RNA virosphere^4–7^. Low fidelity of replication was implicated in limiting genome size in RNA viruses, compared to DNA-based entities^8–10^. This limitation may have been alleviated in viruses with >20 kb genomes by either genome segmentation such as in the dsRNA order *Reovirales*^11,12^, or by acquisition of a proofreading exoribonuclease (ExoN) in the single-segment ssRNA+ viruses of the order *Nidovirales*^13–15^. This virus order represents a monophyletic group of animal viruses with unsegmented or bisegmented^5,16^ genomes and an extraordinarily wide range of RNA genome sizes from 12 up to 41 kb^17–19^, although larger nido-like genomes may exist^20,21^. Nidoviruses are the only known RNA viruses with genomes exceeding 30 kb.

Improving understanding of how nidoviruses have evolved and use such exceptionally large RNA genomes provides a unique window into the primordial life^22^. Also, it informs applied research on many pathogens including severe acute respiratory syndrome coronavirus 2 (SARS-CoV-2), which belongs to the nidovirus family *Coronaviridae*^23^ that together with the nidovirus family *Tobaniviridae* include three viruses with the largest known RNA genomes of vertebrate hosts (36.1-36.7 kb)^16,24,25^. Two of these viruses have bisegmented genomes and infect aquatic hosts^5,16,24^. Based on the protein content, the nidovirus genome can be divided, from the 5’-to 3’-end, into three functional regions that predominantly control genome expression, genome replication, and virus dissemination, respectively^26^. This genomic organization and the associate division of labor may facilitate and constrain genome size change. For nidoviruses with a canonical single-segment genome and multiple open reading frames (ORFs), the three regions correspond to overlapping large ORF1a and ORF1b, and an array of multiple ORFs at their 3’ (3’ORFs), respectively. Upon translation of the genomic RNA by the ribosome, two polyproteins are synthesized: pp1a, encoded in ORF1a, and pp1ab that is produced by extending translation into ORF1b by −1 programmed ribosomal frameshifting (−1 PRF)^27–30^. Proteolytic autoprocessing of pp1a/pp1ab produces subunits of the replication-transcription complex (RTC)^31–33^. In contrast, 3’ORFs encode structural and accessory proteins that are expressed from a nested set of 5’-coterminal subgenomic mRNAs whose RTC-mediated synthesis on the anti-genomic template (transcription) is controlled by discontinuous leader and body transcription-regulating sequences (TRS) in most characterized nidoviruses^34–38^. Other nidoviruses may use leaderless transcription^39,40^.

Nidoviruses are distinguished by an array of five universally conserved domains, including four key enzymes of replication: main 3C-like protease (Mpro or 3CLpro) responsible for pp1a/pp1ab autoprocessing; core RTC components including nucleotidyltransferase (NiRAN) that controls the 5’-terminal structure of viral RNAs and is the nidovirus marker domain; RNA-dependent RNA polymerase (RdRp); and Zn-binding domain (ZBD) fused to superfamily 1 helicase (HEL1)^41^. In the RTC, they are assisted by many other viral non-structural protein products of diverse functions and variable conservation. Experimental characterization of SARS-CoV-2 and a few other nidoviruses defines our current understanding of the RTC function and structure^31–33^. Comparative genomics led to the discovery of most replicase functions and places the obtained experimental results within an evolutionary framework. This includes mutation patterns and lineage- and host-specific variations in the RTC subunit composition^41^.

Unlike 128 known nidovirus species expressing replicase proteins from two overlapping ORFs, two known nidoviruses with extra-large RNA genomes have in-frame ORF1a and ORF1b, either as part of a single genome-wide ORF (41.2 kb genome of planarian secretory cell nidovirus (PSCNV))^17^ or separated by a termination codon (35.9 kb genome of Aplysia abyssovirus 1 (AAbV))^18^. PSCNV and AAbV may use either −1 PRF in a non-canonical way or a readthrough mechanism, respectively, to attenuate genomic RNA translation at a position similar to that of the ORF1a/ORF1b junction. This raises the question whether expansion of RNA genomes to very large sizes in nidoviruses is not supported by canonical replicase ORF organization and expression.

To address this question, here we seek to expand the genomic characterization of invertebrate nidoviruses. We apply a Data-Driven Virus Discovery (DDVD) approach^42^ to mine raw sequencing data from the Sequence Read Archive (SRA)^43^ for divergent RNA viral sequences across a wide eukaryotic host spectrum. We analyze 581,629 eukaryotic SRA transcriptome experiments in a sensitive and highly parallelized fashion, as described previously^16,44–46^. Among the discovered set of RdRp-encoding sequences are 116 nidovirus sequences found exclusively in datasets from animals. All of the new genomes reported here encode the five most conserved replicase domains that distinguish nidoviruses from other viruses. The subset of *vertebrate-*associated nidoviruses is described in another study^16^.

### Discovery of giant invertebrate nidoviruses

We report 76 new nidovirus genomes associated with 72 different *invertebrate* host species (Table S1). All new genomes passed a stringent quality control (Fig. S1) and form numerous divergent phylogenetic lineages within the *Nidovirales* (Figure 1a). We tentatively classify them into 18 viral family-like operational taxonomic units (fOTUs), of which 11 are novel, using phylogenetic analysis and DEmARC, a quantitative, RdRp sequence-based classification approach^47,48^ (Fig. S2, Table S2). Notably, only six out of the 76 discovered nidovirus genome sequences show >90% sequence similarity to five partial and one coding-complete nidovirus genomes found in another DDVD study in which SRA data sets were analyzed^5^, and only three similar sequence fragments were reported in another large-scale RNA virus discovery study^6^ (Table S3). These comparisons show that the data analysis pipeline used in our study has excellent sensitivity regarding the detection and genome sequence assembling of novel nidoviruses with remote sequence similarity to known species. Many of the newly discovered nidovirus genomes have never-before-observed features of functional and evolutionary significance.

**Fig. 1.**
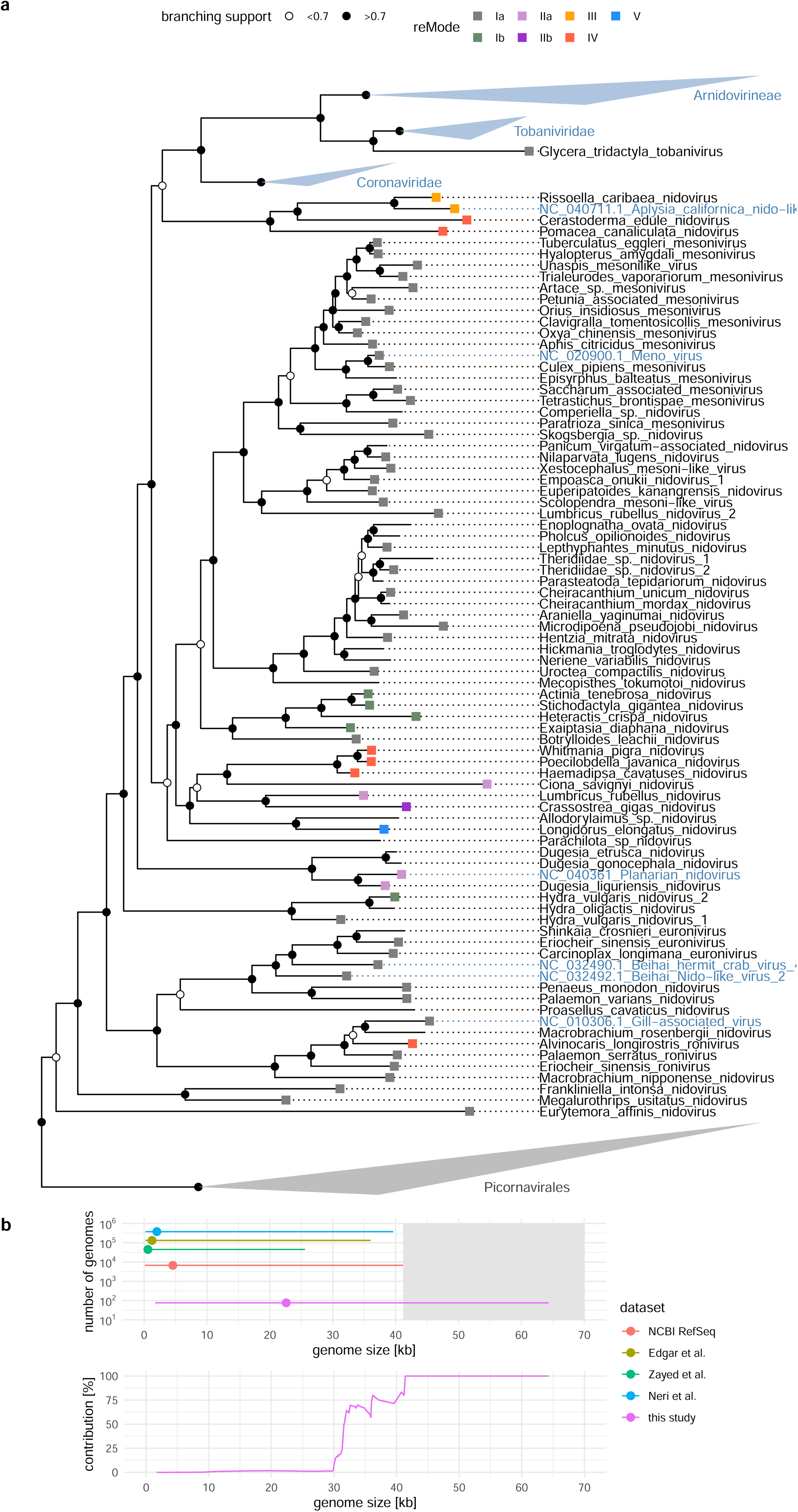
Phylogeny and genome sizes of newly discovered nidoviruses. **a**, RdRp phylogeny reconstructed using PhyML with SH-like branching support. *Picornavirales* reference sequences are used as an outgroup and were collapsed (gray triangle). Names of novel nidoviruses reported in this study are in black and those of selected reference viruses are in blue. White and black circles at internal nodes indicate SH-like branching support smaller and larger than 0.7, respectively. Colored squares at tips indicate the replicase expression mode (reMode). Major vertebrate nidovirus taxa were collapsed (blue triangles). **b**, top: genome sizes ranges across RNA viruses in NCBI RefSeq and three recent large-scale virus discovery studies^5–7^ and nidovirus genome size range reported in this study; circles indicate median values. bottom: percentage of RNA viruses contributed by this study within particular genome size ranges.

Among the 76 discovered invertebrate nidoviruses are 41 with coding-complete genomes (as well as three which we consider nearly coding-complete) of sizes between 18.1 to 64.3 kb (Table S1, Fig. S3). These viruses account for over 50% of ICTV-recognized RNA virus species with genomes larger than 32 kb and 100% of genomes larger than 41.2 kb (Fig. 1b). The newly discovered nidoviruses include viruses with either unsegmented or segmented genomes. The genomes of Crassostrea gigas nidovirus (CGNV) from an oyster, Pomacea canaliculata nidovirus (PCNV) from a snail and Poecilobdella javanica nidovirus (PJNV) from a leech have lengths of, respectively, 64.3, 54.1 and 45.8 kb, making them the largest known RNA viruses together with a 47.3 kb nido-like viral genome reported recently^20^. These viral genomes exceed the median RNA virus genome size of 10-11 kb (such as of hepatitis C virus or human immunodeficiency virus 1) four-to six-fold (Figs. 1b and 2).

**Figure 2.**
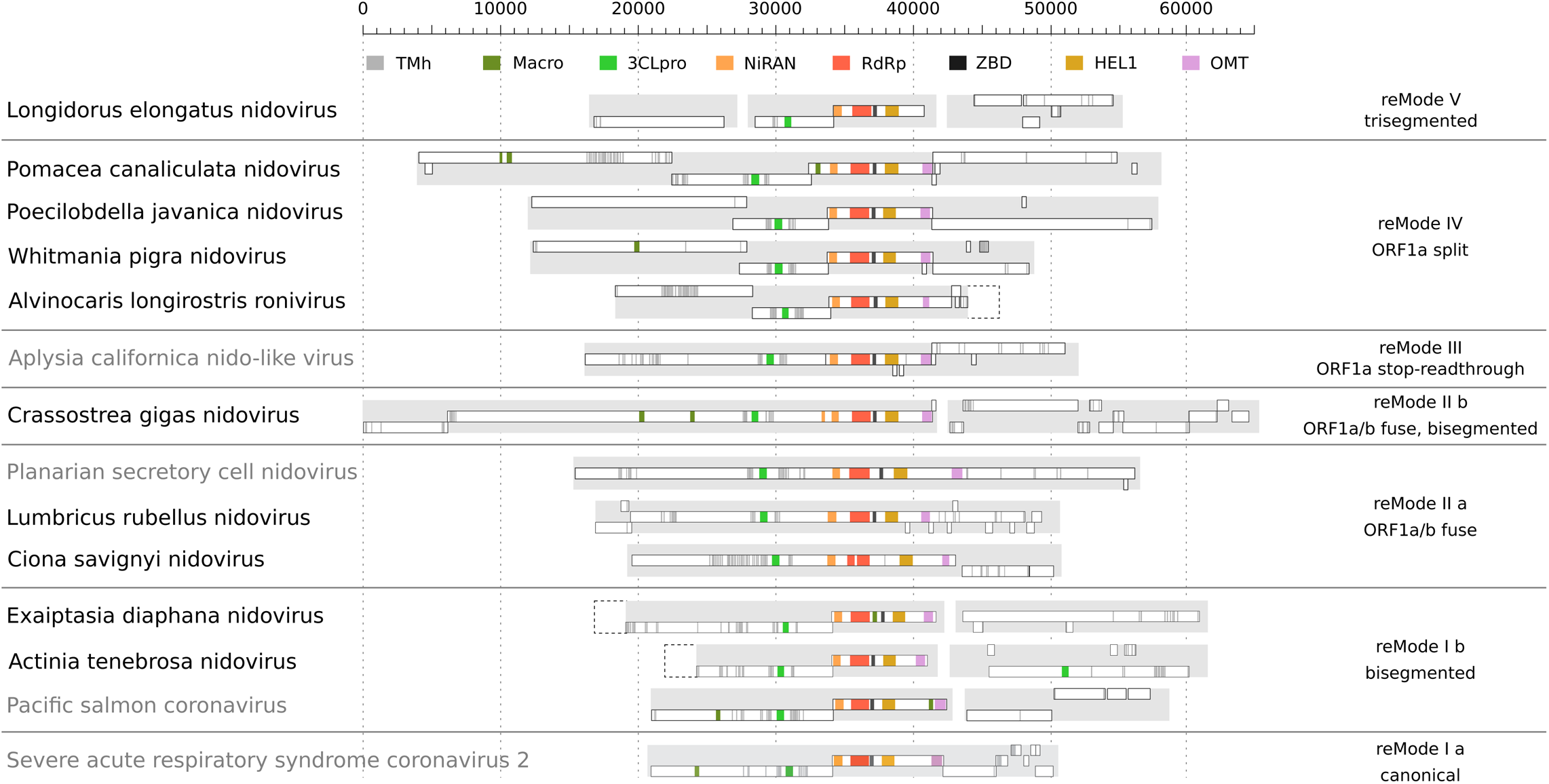
Genome size, polyprotein size and mechanisms of regulating translation of replicative proteins in nidoviruses. **a**, Genome size distribution of novel nidoviruse reported in this study (yellow), reference nidoviruses (blue) and all other reference RNA viruses from NCBI/RefSeq (gray). **b**, same as a but showing length of the longest encoded polyprotein or protein. **c**, Genomic organizations of selected nidoviruses with different replicase expression modes (reModes); drawn to scale in units of nucleotides. Five main modes and four submodes are indicated to the right. Novel nidoviruses and reference viruses have black and gray names, respectively. White rectangles indicate predicted ORFs of at least 300 nt in length delimited by termination codons at both ends. Significant sequence homology to profile Hidden Markov Models of key replicative nidovirus proteins are shown in color. Rectangles with dashed borders indicated sequence of unknown length predicted to be missing in incomplete assemblies. The reference viruses are based on the following NCBI Genbank/RefSeq entries: Aplysia californica nido-like virus (NC_040711.1), Planarian secretory cell nidovirus (NC_040361.1), Pacific salmon coronavirus (MK611985.1), Severe acute respiratory syndrome coronavirus 2 (NC_045512.2).

### Classification of nidoviruses based on replicase genomic organizations

The 76 discovered viruses have diverse genome coding organizations that implicate different mechanisms for the expression of the RdRp and other RTC components. Based on the number of genome segments and organization of ORF(s) encoding the replicase proteins, we recognized five (plausible) modes of replicase expression (denoted ‘reModes’ from I to V) and four submodes of two reModes that are denoted with suffix ‘a’ or ‘b’ after the respective reMode literal (Figs. 1a and 2). This classification is defined by variation at three genome regions listed from 5’-end to 3’-end and including key signals of genome expression: upstream of 3CLpro (the N-terminal most cognate 3CLpro cleavage site), upstream of NiRAN (ORF1a/b −1PRF), and downstream of OMT (ORF1b terminal codon) in coronaviruses. They demark beginning of 3CLpro control over replicase pp1a/pp1ab processing, separate loci encoding accessory subunits and key replicase enzymes of the RTC, and separate replicase from structural genes, respectively. This genome organization and expression may be considered canonical. Three of the reModes accommodate already described mechanisms. The first includes two submodes, the canonical ORF1a/b −1PRF-based mechanism of unsegmented genomes (reMode Ia; n=40 newly discovered virus species) and bisegmented genomes that encode replicase and structural genes on separate segments (reMode Ib; n=5). The two others are employed by a single-ORF PSCNV (reMode IIa; n=3) and AAbV whose ORF1a and ORF1b are fused by a readthrough stop codon (reMode III; n=1) (Fig. 1a and Fig. 2). The remaining two reModes and one sub-reMode are novel and are characterized by, respectively, two –1PRF that elongate translation of the viral genome RNA at junctions in two pairs of overlapping ORFs (reMode IV; n=6), tripartite genome segmentation including one affecting replicase coding and expression (reMode V; n=1), and two –1PRF the second of which attenuates translation of a single ORF (in a manner described for PSCNV) encoded on segment 1 of the bisegmented CGNV (reMode IIb; n=1) (Fig. 2).

Nidoviruses of invertebrate hosts using canonical reMode 1a greatly outnumber all others combined at the species(-like) rank (78 vs. 19), but not at the family(-like) rank (10 vs. 9) (Table S3), although this sub-mode remains the best sampled (Fig. 3a). Thus, invertebrate nidoviruses with non-canonical reModes (Ib-to-V) may be much more common than suggested by their current low sampling. Indeed, majority of virus species using reMode Ia cluster separately from others in few family-restricted monophyletic lineages (Fig. 1a). Likewise, group-specific phylogenetic clustering also is observed for viruses of reModes Ib, IIa, III, and IV, which include two or more viruses (Fig. 1a). It implies that once emerged, a reMode-specific genomic make-up persists in evolution. Furthermore, each of reModes Ib, IIa, and IV are observed in two very distantly related lineages, indicating that they may have emerged independently at least twice during the evolution of nidoviruses. In most instances, viruses of reModes Ib-V are sister to canonical nidoviruses of reMode Ia, which thus may represent the ancestral state for others. Based on tree topology, this seems to be most likely for two groups of viruses using reMode Ib and Alvinocaris longirostris ronivirus (ALRV), a shrimp virus of reMode IV that is part of the otherwise reMode Ia clade representing the family *Roniviridae*. There are two exceptions: two pairs of nidoviruses of reModes III and IV that were derived from mollusk projects, which are sister to each other and are deeply rooted in the nidovirus phylogeny (Fig. 1a, Fig. 2, Table S2). A much larger sampling may be required to reconstruct the full evolutionary history of reModes.

**Figure 3.**
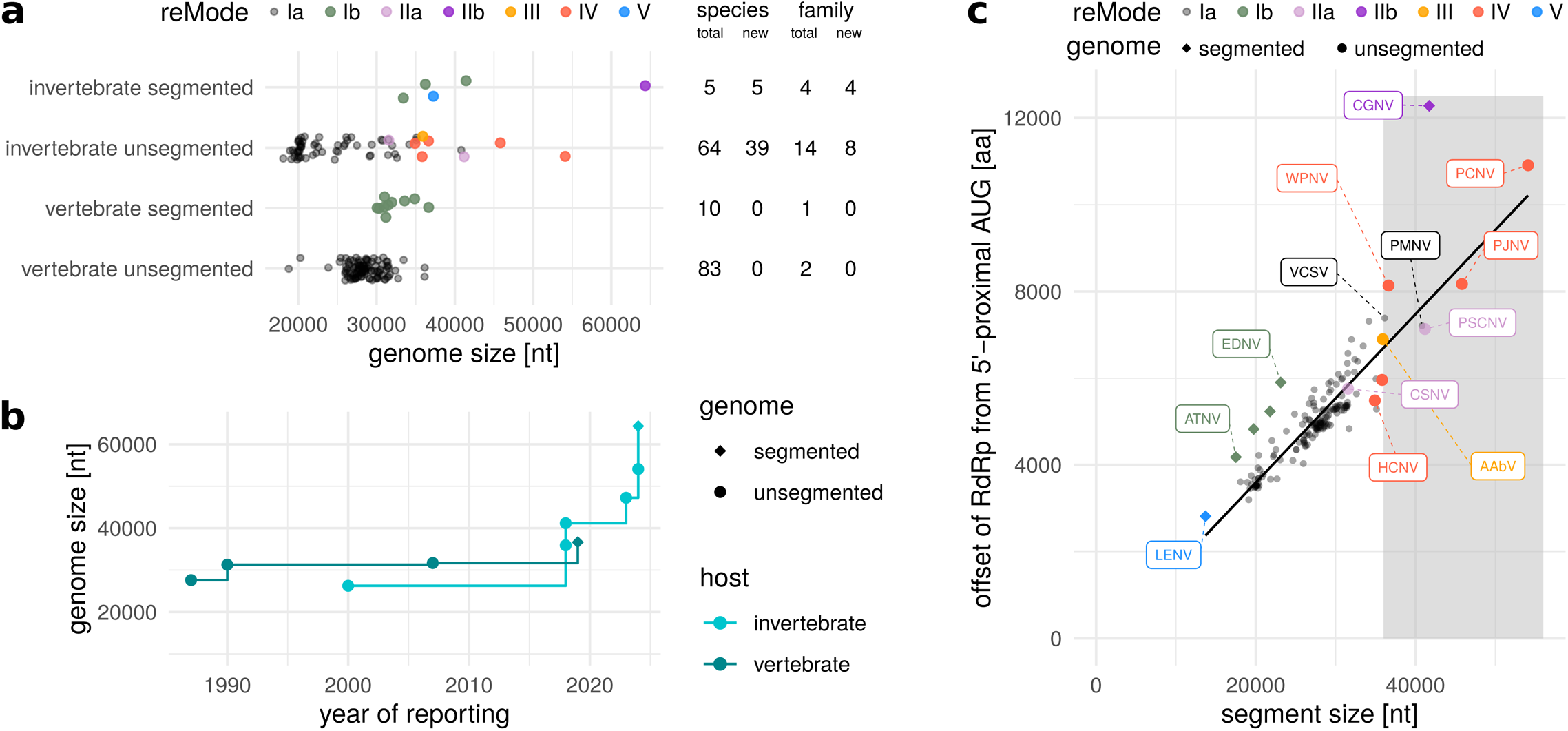
Nidovirus genome size expansion. **a**, Genome sizes of ExoN-encoding vertebrate and invertebrate nidoviruses with segmented and unsegmented genomes. Only coding-complete genomes are considered. Point color indicates replicase expression mode (reMode). The number of total and new virus species and virus families per group is indicated at the right; note that a virus family-like OTU may include both viruses with segmented and viruses with unsegmented genomes. **b**, Size increase of largest known genome of vertebrate and invertebrate nidoviruses from 1987 until 2024. **c**, Relation of the length of the largest segment (which corresponds to genome size for unsegmented viruses) and the position of the RdRp in the pp1ab polyprotein. Only newly discovered nidoviruses with coding complete genomes are included in addition to 156 coding-complete nidovirus genomes from NCBI RefSeq. Viruses with segmented und unsegmented genomes are shown as diamonds and circles, respectively. Symbol color indicates reMode. The gray background highlights the genome size region above 36 kb. The following viruses are labeled: Longidorus elongatus nidovirus (LENV), Actinia tenebrosa nidovirus (ATNV), Exaiptasia diaphana nidovirus (EDNV), Whitmania pigra nidovirus (WPNV), Pomacea canaliculata nidovirus (PCNV), Poecilobdella javanica nidovirus (PJNV), Haemadipsa cavatuses nidovirus (HCNV), Ciona savignyi nidovirus (CSNV), Crassostrea gigas nidovirus (CGNV), Penaeus monodon nidovirus (PMNV), Veiled chameleon serpentovirus (VCSV, NC_076911.1), Planarian secretory cell nidovirus (PSCNV, NC_040361.1), Aplysia californica nido-like virus (AAbV, NC_040711.1).

### Association of replicase expression modes with genome expansion trajectories

ExoN-encoding viruses of canonical reMode I belong to two families of vertebrate nidoviruses and at least nine families or family-like OTUs of invertebrate nidoviruses. They have genomes in a size range from 20 to 36 kb (Fig. 3a). For vertebrate nidoviruses, a genome size of 36 kb seems to be close to the upper limit, given how modestly (9 kb) the largest genome size of vertebrate-associated nidoviruses increased over 38 years since the first coronavirus genome was reported (Fig. 3b). Their counterparts of four other reModes (II-V) belong to seven families or family-like OTUs of invertebrate nidoviruses (including ALRV) with 31-to-64 kb genomes. A relatively small overlap of 5 kb between the two genome size ranges (Fig. 3a) indicates a size threshold in the expansion of RNA genomes that interconnects vertebrate/invertebrate host divide and diversification of reModes. Notably, for both vertebrate and invertebrate nidoviruses, the largest genome is segmented (Fig. 3b) with segment 1 coding for the replicative proteins and segment 2 for the structural and accessory proteins. Also the other nidoviruses with bisegmented (reMode Ib) and trisegmented (reMode V) genomes have sizes in the upper range for viruses of their respective host group (Fig. 3b), suggesting that genome segmentation is one possible pathway to evolve very large RNA genomes (similar to the multi-segment genomes in the size range from 18 to 30 kb in the order *Reovirales*^11,12^). Interestingly, none of the segmented nidovirus genomes of either veretebrate or invertebrate hosts are smaller than 30 kb, indicative of a nidovirus-specific constraint. Alternative genome expansion pathways are supported by the other non-canonical reModes, as almost all viruses with genomes larger than 36 kb are either segmented (sub-reMode Ib) or of reModes II-V, with only three reMode Ia exceptions: the 36.1 kb Veiled chameleon serpentovirus^25^, the 36.1 kb Hypomesus transpacificus coronavirus^16^ and the 40.8 kb Penaeus monodon nidovirus from this study (Table S1).

A strong linear correlation of the genomic position of the RdRp gene with genome/segment size (Fig. 3c) demonstrates that a size increase of the ORF1a(-like) region in ancestral viruses was a primary factor in the emergence of the nidovirus lineages with giant genomes. This expansion of the ORF1a-like region is most pronounced in CGNV, although segment 2 encoding putative structural proteins of this virus is also the largest among all nidoviruses (Fig. 2). This observation is compatible with a wave-like expansion model that implies varying contributions of specific genomic regions to genome size increase at different points on the genome expansion trajectory^26^.

### Dual ribosomal frameshifting in a subset of nidoviruses with giant genomes

In reMode IV viruses, which include PCNV, PJNV, and four other invertebrate viruses in three distinct clades in the nidovirus tree (Fig. 1a), this region-specific expansion was associated with recurrent replacement of the exceptionally large ORF1a region by two overlapping ORFs, denoted ORF1a1 and ORF1a2 from here on, with the overlap locating upstream of 3CLpro (Figs. 1a and 2). Otherwise, reMode IV resembles reMode Ia. Notably, this genomic organization implies that translation of the RdRp locus in these viruses from the incoming genomic RNA must involve two instead of one –1PRF event in reMode Ia, as this essential protein is synthesized early in the viral replication cycle. In addition, such mechanism would constitute an additional layer of controlling the stoichiometries of viral replicase proteins encoded in ORF1a1 relative to those in ORF1a2 and ORF1b.. In coronaviruses, a spatially equivalent genomic region encodes multiple functions including subunits of a molecular pore that spans the double membrane of the coronavirus replication organelle^49^.

The presence of bioinformatically predicted frameshift signals in and immediately downstream of the ORF1a1-ORF1a2 and the ORF1a2-ORF1b overlapping region in the PCNV genome, including pseudoknot RNA structures and identical slippery sequences (UUUAAAC), strongly support this novel reMode IV expression mechanism. Similar genetic elements with PRF signals were identified in the ORF-overlapping regions of most of the other reMode IV viruses (Fig. S4).

To provide experimental evidence for this mode of RNA translation and to check whether the putative frameshift sites are functional in cells, we have employed the dual fluorescence reporter assay that we developed previously^50,51^. To generate dual-fluorescence reporter constructs approximately 150 nt fragment of each nidoviral genome spanning the putative programmed ribosome frameshift sites were placed between the coding sequence of EGFP and mCherry (see Methods for details, Fig. 4a,b). Accordingly, we calculated the efficiency of frameshifting in both −1PRF sites separately in CGNV, WPNV, PCNV, and HCNV. Both sites in CGNV, RFS1 and RFS2, were leading to 0.03 and 2.3% frameshift, respectively. In the other viral sequences significant levels of frameshift were observed at least in one of the frameshift sites (Fig. 4c). Although RFS1 of WPNV only had 6.6% frameshift, RFS2 was much more efficient with 39.9%. PCNV RFS1 and RFS2 had reasonable frameshift levels ranging between 23.1 and 11.5%, respectively. RFS1 of HCNV was only 1.53% efficient, yet the highest level of frameshifting was measured at the HCNV RFS2 by 83%, which is among the highest *in vivo* measured frameshift efficiencies reported so far in viral sequences (Fig. 4c). Of note, the amount of frameshift varied widely among the different viral mRNAs. This aligns with the notion that the pseudoknot structure directly flanking the slippery site not only contributes to increased levels of frameshifting, but plays a direct role in fine tuning the exact amount of −1PRF needed by the virus to regulate the production of viral proteins at the correct stoichiometry from each reading frame. In vivo frameshifting may be further affected by host and virus factors not available in the tested systems.

**Figure 4.**
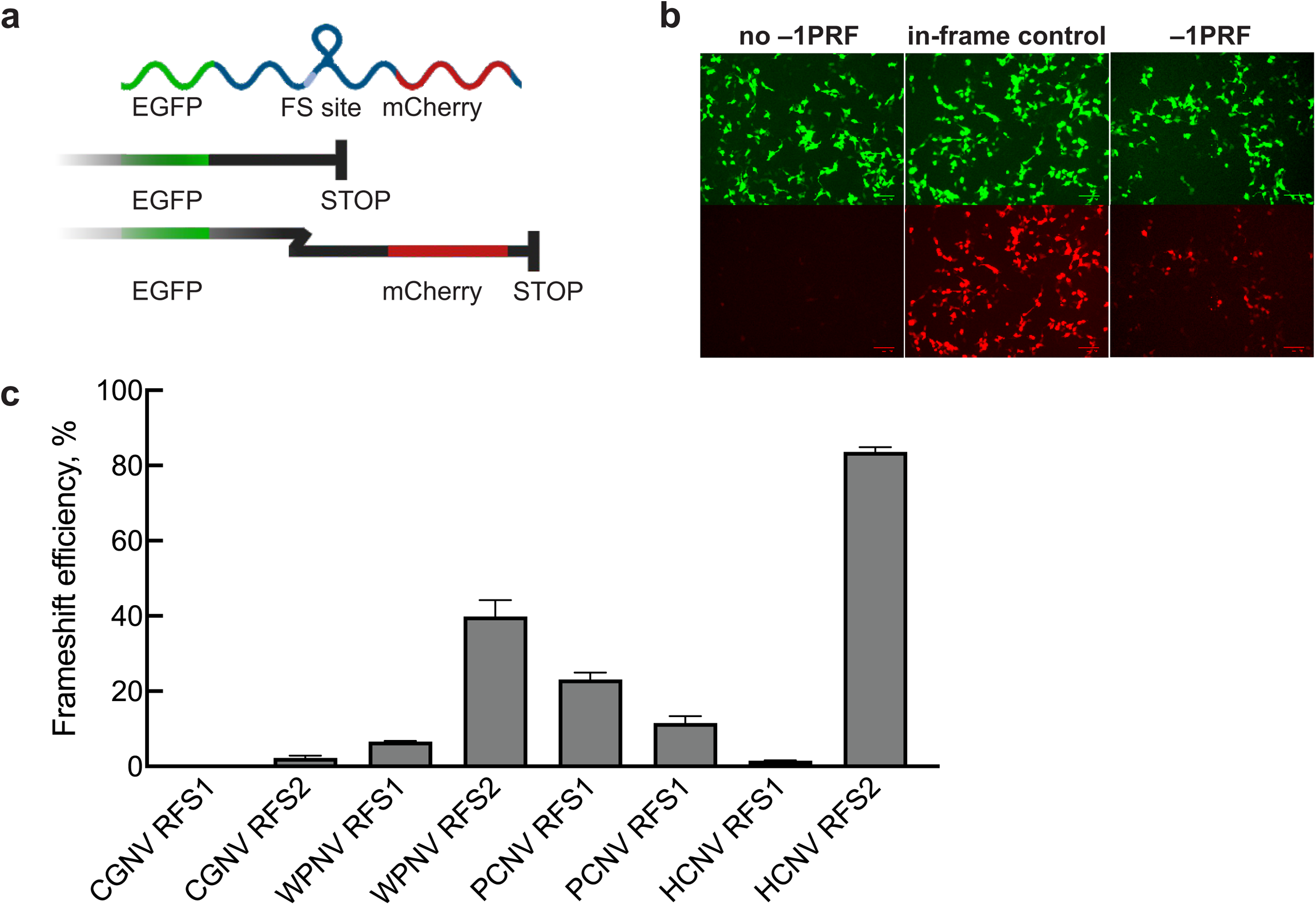
Measuring dual ribosomal frameshifting efficiency in four nidoviruses. **a**, Schematic representation of the dual-fluorescence frameshift reporter construct. EGFP and mCherry are separated by a self-cleaving 2LA peptide as well as by a stop codon in-frame with EGFP. As a result, 0-frame translation would produce only EGFP, whereas –1PRF would produce both EGFP and mCherry. The ratio of mCherry to EGFP fluorescence is used to quantify the FE. **b**, Confocal microscopy images of cells transfected with the EGFP-mCherry control construct lacking frameshift signal (no –1PRF), an in frame control and –1PRF constructs. **c**, Comparison of relative frameshift efficiency of different nidovirus frameshift sites. Data points represent the meanL±Ls.d. (nL=L3 independent experiments). Frameshift efficiency was measured for two ribosomal frameshift sites (RFS) in Crassostrea gigas nidovirus (CGNV), Whitmania pigra nidovirus (WPNV), Pomacea canaliculata nidovirus (PCNV) and Haemadipsa cavatuses nidovirus (HCNV).

The dual frameshifting mechanism in the reMode IV viruses results in elongation of translation of the viral genomic RNA upon –1PRF at both sites to express the RdRp and the other replicase proteins. A similar two-site mechanism is employed by the sole member of reMode IIb viruses, CGNV. Here, −1 PRF at the first site located within the overlap of the two ORFs encoded by segment 1 results in translation progression, while –1PRF at the second site located in the N-terminal NiRAN coding region attenuates translation (Figure 2, Fig. S4). This shifting of ribosomes out of the coding reading frame at the second CGNV site to regulate the stoichiometries of replicase and other non-structural proteins resembles the mechanism described for PSCNV^17^. Compared to all other nidoviruses, CGNV is also exceptional with respect to the position of the RdRp (and other replicase proteins) in the synthesized polyprotein being more than 12000 amino acids C-terminal of the polyprotein N-terminus. These characteristics, together with the bisegmentation of its giant 64.3 kb genome and possible production of subgenomic RNAs from segment 2, demonstrate that the control of gene expression in CGNV is one of the most sophisticated across the RNA viruses.

### The first nidovirus with a trisegmented genome

The sole member of eType V viruses, Longidorus elongatus nidovirus (LENV), has a trisegmented 37.3 kb genome which encodes ORF1a1, ORF1a2/ORF1b, and the structural ORFs on different segments (Fig. 2). The consistently high read coverage depth values, which well exceed 100 reads per genomic positions for all three segments (see next section for more details), strongly suggest that virus origin by contamination of the specimen is unlikely. Conserved sequence termini that are shared among the three segments and which end with expected poly(A) tails in case of the 3’-ends show that the trisegmentation of the genome is authentic (Fig. S5), making LENV the first known nidovirus with three genome segments. This trisegmented genomic organization enables differential regulation of viral protein amounts via segment-specific copy numbers.

### Transcriptional analysis of non-canonical reMode viruses

SRA datasets containing novel nidoviruses were further examined for three types of RNA data commonly associated with subgenomic transcription in nidoviruses: a) presence of putative leader and body transcription regulatory sequences (TRS) upstream of important structural genes, b) stepwise increases in sequence read depth associated with production of one or more 3’-coterminal sgRNA species, and c) reads joining the genomic leader sequence to a downstream body sequence at a TRS. Whitmania pigra nidovirus (WPNV) (reMode IV) and Ciona savignyi nidovirus (CSNV) (reMode IIa) both show polar read depth increases as well as presence of putative leader-body-junction reads consistent with canonical nidovirus subgenomic RNA production (Fig. S6). In contrast, CGNV (reMode II b) shows stepwise read depth increases at two putative TRS, but does not contain any sequences that would be consistent with leader addition (Fig. 5). CGNV may therefore produce leaderless sgRNA species, as reported for two other previously characterized invertebrate nidoviruses, GAV^52^ and AAbV^39^, but with leader TRS at the 5’-end of segment 1, and body TRS on segment 2 (Fig. 5). LENV (reMode V) does not show marked read depth increases at putative TRS upstream of three inferred structural genes on segment 3. Notably, and uniquely among all known multi-segment nidoviruses, all three segments of LENV contained a near-identical 206 nt sequence at the 5’-termini (Fig. S5), suggestive of a new expression strategy that may be related to canonical nidovirus discontinuous transcription, leader-switching^53–55^ or both (Fig. 5). Taken together with results for reMode I viruses^52,56^ and reMode III^39^, there is evidence for production of 3’-coterminal sgRNA species, a hallmark of canonical nidovirus replication, across all nidovirus reModes.

**Figure 5.**
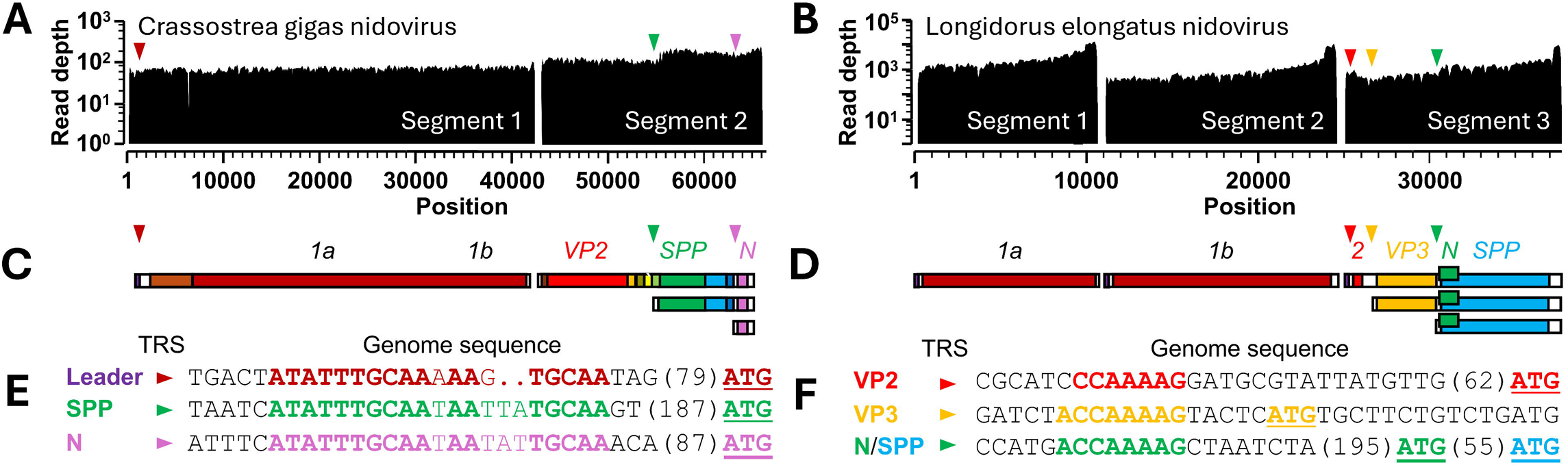
Evidence for subgenomic RNA production in two segmented nidoviruses. Sequence read depth maps (A, B), maps of genome coding capacity (C, D), and putative transcription-regulatory sequences (E-F) are shown for Crassostrea gigas nidovirus (reMode IIb; panels A, C, E) and Longidorus elongatus nidovirus (reMode V; panels B, D, F). Inferred start codons are underlined and color-coded to match the genome maps. VP2, virus protein 2; SPP, structural polyprotein with similarity to that of alphaviruses; N, nucleoprotein-like basic soluble protein.

## Conclusions

In summary, our study employing a Data-Driven Virus Discovery approach reveals several novel evolutionary trajectories taken by different nidoviruses during evolution of the largest and most complex RNA genomes known. Nidoviruses therefore provide a unique window into the evolution and function of RNA genomes of sizes that are not accessible to other viruses. In all known nidoviruses using proofreading ExoN, we document restricted RNA genome size ranges that are associated with genome segmentation, genomic organization, or infected animal host. The genome size range of invertebrate nidoviruses is more than twice as large as that of vertebrate nidoviruses. In the current dataset, invertebrates are represented by only a subset of known phyla, which may be a more informative host denominator to use with improved nidovirus sampling. Multiple nidovirus lineages have independently reached states of high genetic complexity by evolving exceptionally large genomes and sophisticated translational control mechanisms such as dual frameshifting. The function of dual frameshifting in the nidovirus life cycle, the influence of body temperature of invertebrate hosts on frameshifting efficiency, how cis-acting RNA elements lead to such varying levels of frameshifting and whether these sites are additionally regulated by host and viral trans-acting factors will remain to be explored. Overall, our findings pose the question whether similar mechanisms are employed by other viruses.

## Methods

### SRA-based virus discovery

Our computational virus discovery workflow and its application to SRA data is described in detail in a previous study^16^. It is highly parallelized and was run on a high-performance computing cluster. The workflow involves sensitive detection of raw viral sequencing reads matching to nidovirus NiRAN or RdRp protein profile Hidden Markov Models using hmmsearch of the HMMer package^57^ and targeted (seed-based) viral genome assembly using Genseed-HMM^58^ applied to all sequencing experiments identified in the first step. Seed-based nidoviral genome assemblies were validated and, in some cases, extended using untargeted de novo assembly using Megahit^59^ and Spades^60^, and manually curated. Adapter sequences and low-quality bases were trimmed using fastp^61^ prior to viral genome assembly.

For viruses with segmented genomes, additional segments were identified by (i) searching for significant sequence similarity, via a nucleotide-based Blast search, between the termini of the identified RdRp-encoding segment of a virus and those of other contigs derived from the same dataset, and/or (ii) searching for significant sequence similarity, via a protein-based Blast search, between structural nidovirus reference proteins and contigs of the dataset from which the RdRp-encoding segment was derived. For Crassostrea gigas nidovirus (CGNV), which encodes the RdRp and other replicase proteins at the 3’-proximal part of its 41.7 kb segment 1 (Figure 2), the two methods did not yield a potential second segment. Therefore, we applied the following strategy. We considered all 626,006 contigs assembled *de novo* from this SRA dataset (accession SRR15293269) and kept those that (i) have an average coverage depth larger than the average coverage depth of CGNV segment 1 (559.02) and (ii) that are at least 4 kb in length, leaving 24 contigs. We further filtered the contigs by only keeping those for which at least 50% of the sequence was covered by ORFs predicted using EMBOSS getorf^62^ from which at least one ORF was longer than 2 kb, leaving 9 contigs. We reasoned that the major part (virtually everything except the terminal untranslated regions) of an RNA viral sequence is protein-coding. We then conducted a Blastx^63^ search of these 9 contig sequences against Genbank nr_clustered and discarded those that gave strong hits against host proteins with an E-value of 1e-10 or better and a sequence identity of at least 50%, leaving three contigs. The first one was CGNV segment 1. The second contig was a ∼9 kb coding-complete picornavirus-like genome sequence that showed about 70% amino acid sequence identity to Beihai picorna-like virus 107 (NC_032617). The third contig, deemed CGNV segment 2, was ∼22.6 kb in length, had a poly-A tail of length 53, and coded for 10 ORFs (>450 nt) that covered almost the entire sequence, the largest of which was >8 kb. It was the second largest contig in the whole dataset (after CGNV segment 1) and showed a mean coverage depth of 1972.2. Only a very short region of CGNV segment 2 (<2% of its full length) gave hits in the Blastx search again Genbank nr_clustered with <30% amino acid sequence identity to a host urokinase protein, a serine protease. The remaining part of the CGNV segment 2 sequence did not receive any hits in this Blastx search. However, eight of the ten CGNV segment 2-encoded ORFs gave hits (probabilities 30.6-96.2%) in a HHpred^64^ search against different viral proteins, including a hit against a viral Fusion glycoprotein (PF13044.10, Table S4), but not against viral replicative proteins.

### Assembly quality assessment

We conducted a systematic quality assessment of each assembled nidovirus genome, as described previously^16^. Briefly, we calculate two metrics per assembled sequence that quantitatively assess the per-base (minimal coverage [mico]) and contig-wide (mean alignment score [meas]) accuracy, respectively. The mico and meas value of each assembled nidovirus sequence is then assessed relative to a reference assembly set composed of RNA virus genome sequences from two conventional virus discovery studies^65,66^. This allows partitioning the mico and meas values into deciles with lower, medium and higher deciles indicating, respectively, lower, average and higher assembly quality.

### Comparison with other large-scale RNA virus discovery studies

We compared our nidovirus genome sequences against RNA viral sequences reported in Edgar et al.^5^ and Neri et al.^6^. To this end we downloaded all RdRp-containing contigs from these studies and used them as a database for a nucleotide Blast with word size of 7 against the nidovirus contigs described in the current study. Only hits with an alignment length and a sequence identity of at least 100 nucleotides and 90 percent, respectively, were considered. In total, only nine of our contigs obtained hits against sequence fragments from the other studies, all except one of which very (much) shorter than the viral sequences reported here (see Table S3 for details). Notably, 48 of the SRA runs in which we identified the novel invertebrate viruses (with highest mean viral sequencing depth) (Table S1) were also analyzed by Edgar et al.^5^ as these sequencing libraries are part of their rindex.tsv (downloaded 6 Sep 2022) of completed RdRp SRA accessions.

### Prediction of ribosomal frameshifting sites

The available full-length genomic sequence of a virus was used as input for slippery sequence and RNA pseudoknot prediction using KnotInFrame^67^. Only hits within the ORF1a1-ORF1a2 or ORF1a2-ORF1b overlapping region, that is from the stop codon delimiting the right-hand ORF at the 5’ side until the stop codon delimiting the left-hand ORF at the 3’ side, were considered. The sequence and predicted structure of RFS elements was visualized using VARNA v3.93^68^.

### Phylogenetic analysis

Multiple RdRp protein sequence alignments of ingroup (nidovirus and nidovirus-like) and outgroup (picornavirus) sequences were computed separately using MAFFT v7.310^69^ with options ‘--localpair -- maxiterate 1000 --retree 3’. To build the outgroup, we retrieved all *Picornavirales* RdRp-containing polyprotein sequences available in NCBI RefSeq in May 2022 and clustered them at 65% protein sequence identity. In- and outgroup alignments were combined using MUSCLE v3.8.31^70^ in profile mode, followed by manual curation. To obtain an alignment derivative of most conserved RdRp regions for phylogenetic analysis, we then excluded, in the following order, alignment columns which had a gap in more than 5% of the sequences and sequences with more than 50% of gaps across all remaining alignment columns, the latter representing sequence fragments from incomplete assemblies. The best-fitting substitution model, which was found to be LG+G4+I, was selected using ModelTest-NG v0.1.7^71^. Phylogenetic reconstruction was performed using PhyML v20120412, RAxML v8.2.12 and BEAST v1.8.0^72–74^. The three tools generated trees with very similar topologies and the PhyML version was used for subsequent analyses and visualization.

### Sequence-based virus classification

We used DEmARC^47,48^ v1.4 to classify viruses into operational taxonomic units (OTUs) at the family and genus level. Briefly, DEmARC implements a pairwise-distance based approach that proposes thresholds on pairwise genetic divergence to group similar viruses into clusters whose members show genetic distances that are predominantly smaller than the chosen threshold. Optimal thresholds are found in a data-driven way by minimizing clustering cost and maximizing clustering persistence. The clustering cost is proportional to the number of intra-cluster distances exceeding a threshold and clustering persistence reflects the range of pairwise distances within which the clustering is stable. We used patristic distances extracted from the nidovirus RdRp phylogeny reconstructed in this study as input for DEmARC. This classification closely reproduces the available classification for nidoviruses at taxonomy ranks above species, which we developed by applying DEmARC to five most conserved replicase domains, including the RdRp^19,23,75^. Since most newly described nidoviruses prototype new species, the reported demarcation at this rank for new nidoviruses is trivial and accurate. For other nidoviruses, we used formally recognized species. We used species taxa^76^ to correct for highly uneven sampling - from singletons to millions of genome sequences per species – in our comparative analyses concerning host, genome size and genome organization.

### Transcriptional analysis

Read depth analysis and BLAST leader-body junction detection were performed as previously described^16,18^ using bowtie2 2.5.1^77^, samtools 1.17^78^, bcftools 1.17^78^ and the NCBI SRA toolkit 2.4.10^43^.

### Assessment of ribosomal frameshifting in cells using dual fluorescence reporters

To generate dual-fluorescence reporter constructs to assess frameshift efficiencies, programmed frameshift sites of each nidovirus were placed between the coding sequence of EGFP and mCherry (parental Addgene plasmid # 8780362) by Gibson assembly such that EGFP would be produced in 0-frame and mCherry in –1-frame with the −1PRF sequence^50,51^. An in-frame 100% mCherry control for each vector was generated to normalize the mCherry fluorescence signal. HEK293 cells were maintained in DMEM (Gibco) supplemented with 10% FBS (Gibco) and 100 μg/ml streptomycin and 100 U/ml penicillin. Cell lines were kept at 37 °C with 5% CO_2_. Transfections were performed with 0.05 pmol of plasmid DNA and PEI (Polysciences) according to manufacturer’s instructions. Cells were harvested at 24 h post-transfection and gently washed with PBS for flow cytometry. Flow cytometry was performed on a NovoCyte Quanteon (ACEA) instrument and data were analyzed with FlowJo software (BD Biosciences). Frameshifting efficiency (FE) was calculated according to:

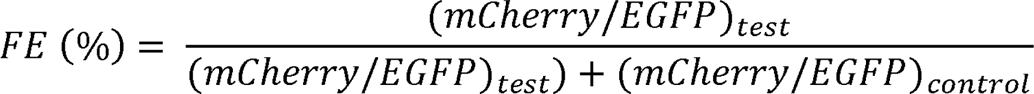

where mCherry represents the mean mCherry intensity, EGFP the mean EGFP intensity, test represent the tested sample and control represents the in-frame control where mCherry and EGFP are produced in the same frame at an equimolar ratio. Data represent the results of at least three independent experiments.

## Data availability

All data used in this study are publicly available through the Sequence Read Archive repository. Viral genome sequences assembled in this study will be made available through the NCBI Genbank database upon publication.

## Code availability

The Virushunter and Virusgatherer tools are available on github: https://github.com/lauberlab/VirusHunterGatherer.

## Acknowledgements

C.L. presented the virus discovery part and initial data on dual ribosomal frameshifting reported in this paper at the international Nido2023 symposium (Montreux, Switzerland, May 2023). C.L. is supported by the Deutsche Forschungsgemeinschaft (DFG, German Research Foundation) under Germany’s Excellence Strategy - EXC 2155 - project number 390874280. C.L., S.S. and R.B. acknowledge support of the Project “Virological and immunological determinants of COVID-19 pathogenesis – lessons to get prepared for future pandemics (KA1-Co-02 “CoViPa”)”, a grant from the Helmholtz Association’s Initiative and Network Fund. N.C. is funded by the Helmholtz Association and European Research Council-project number 948636. The funders had no role in study design, data collection and analysis, decision to publish, or preparation of the manuscript. We thank Dmitry Samborskiy (Moscow State University) for help with DEmARC. We thank all colleagues in the scientific community who make their sequencing data publicly accessible. We acknowledge the NCBI for providing an elaborate platform to exchange sequencing data. We thank the Center for Information Services and High-Performance Computing (ZIH) at TU Dresden for generous allocations of computer time. C.L. and A.E.G. are members of the European Virus Bioinformatics Center (EVBC).

## Contributions

C.L., S.S., N.C. and A.E.G. conceptualized the study approach. C.L., N.C. and R.B. supervised the research. A.S. performed in vivo dual frameshift reporter experiments. C.L., A.S., J.V., S.S, B.W.N., and A.E.G. analyzed the data. Data were interpreted by C.L., B.W.N, S.S., N.C. and A.E.G. Data visualizations were created by C.L., A.S. and A.E.G. The paper was drafted by C.L. and A.E.G. and was reviewed, edited and approved by all authors.

## Supplementary Information

**Table S1.** Summary information of the 76 newly discovered nidovirus genomes and SRA datasets they were retrieved from.

**Table S2.** DEmARC-based classification of the discovered nidoviruses.

**Table S3.** Results of blastn search with word size 7 and percent identity and alignment length cut-offs of 90 and 100, respectively, of the 76 discovered nidovirus genome sequences against RdRp-containing contigs reported in Edgar et al. (Ref.5) and Neri et al. (Ref.6).

**Table S4.** Identification of genome segment 2 of Crassostrea gigas nidovirus (CGNV).

**Fig. S1.** Viral genome assembly quality metrics.

**Fig. S2.** DEmARC analysis.

**Fig. S3.** Genomic organization of 76 newly discovered invertebrate nidoviruses.

**Fig. S4.** Prediction of pseudoknot structures and slippery sequences in reMode IV viruses using KnotInFrame.

**Fig. S5.** Conserved sequence at segment termini of LENV. Conserved sequence termini of the three LENV segments. Shown are the 5‘ (A) and 3‘ (B) sequence termini of contigs produced by separate, segment-specific assemblies (top three sequences) and a joint assembly of all three segments (bottom three sequences). A seed-based assembly strategy was used and the coding regions of the segments formed the seeds.

**Fig. S6.** Evidence for subgenomic RNA production in two unsegmented nidoviruses. Sequence read depth maps (A-B), maps of genome coding capacity (C-D), putative transcription-regulatory sequences (E-F) plus sequences and read counts containing subgenomic RNA leader-body junctions (G-H) are shown for Whitmania pigra nidovirus (reMode IV; panels A, C, E, G) and Ciona savignyi nidovirus (reMode II a; panels B, D, F, H). Inferred start codons are underlined and color-coded to match the genome maps. VP2, virus protein 2; SPP, structural polyprotein with similarity to that of alphaviruses; N, nucleoprotein-like basic soluble protein.

