## Supplementary figures and images for "RNA genome expansion up to 64 kb in nidoviruses is host constrained and associated with new modes of replicase expression"

### Figure S1

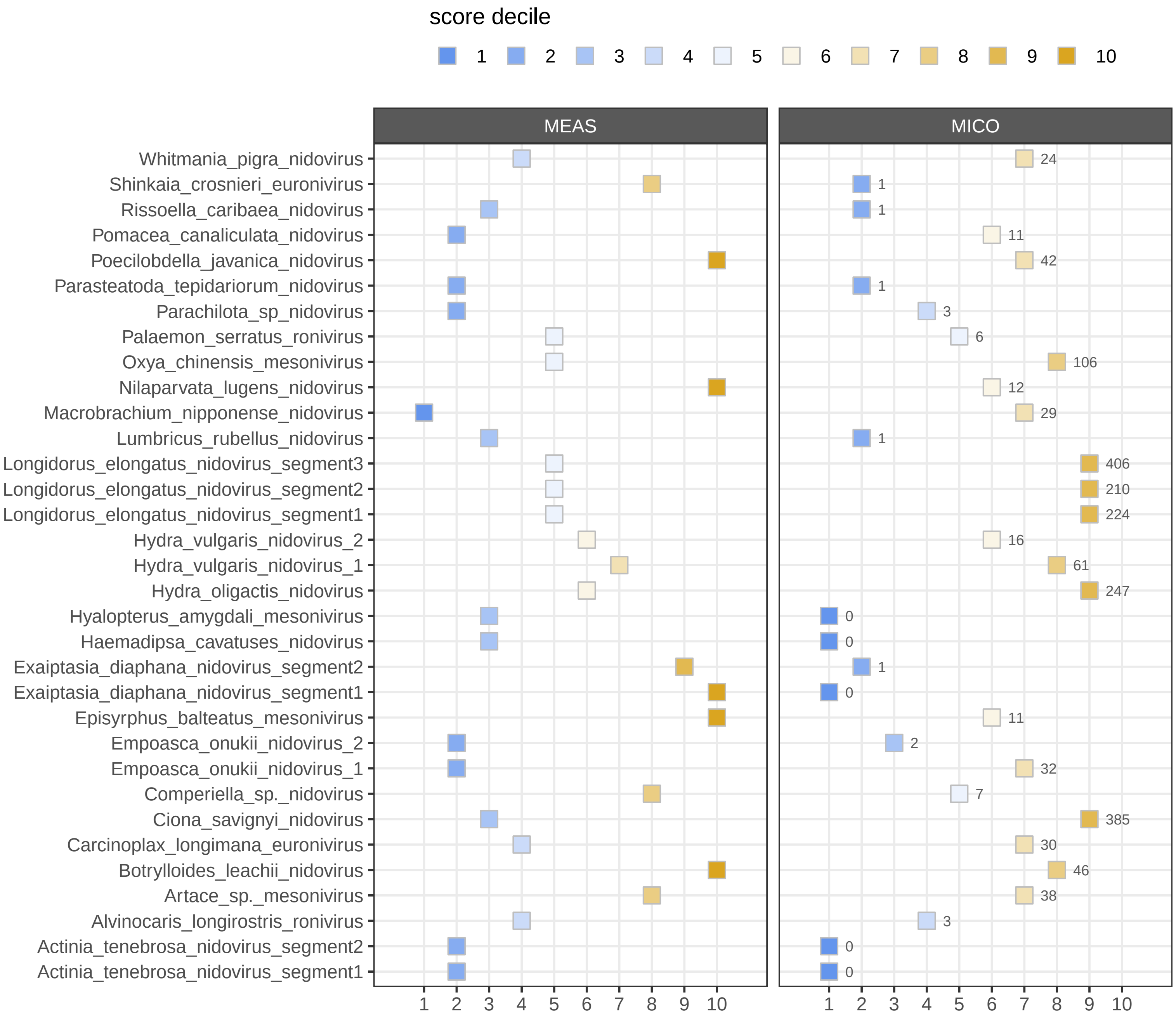

### Figure S2

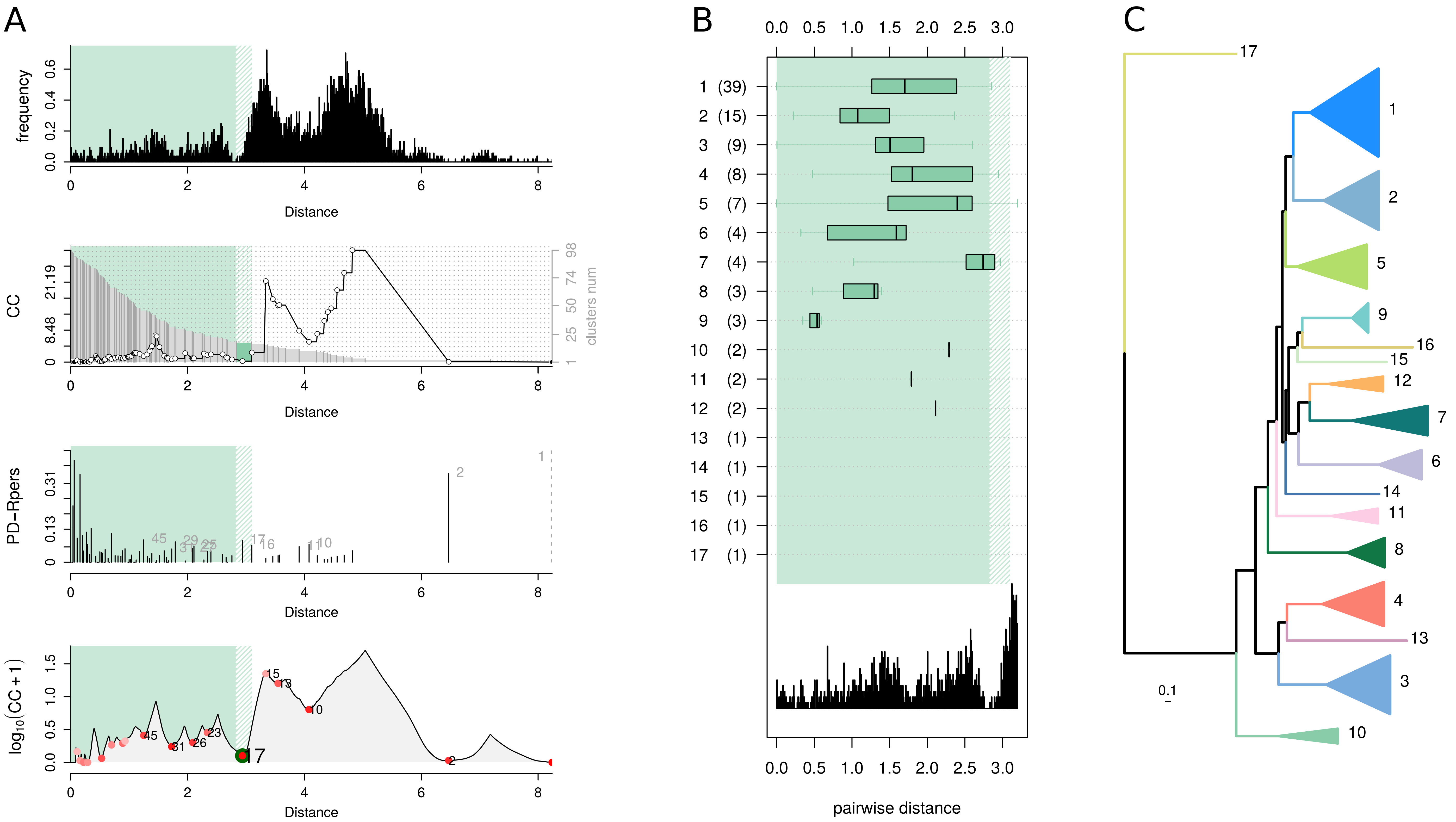

### Figure S3

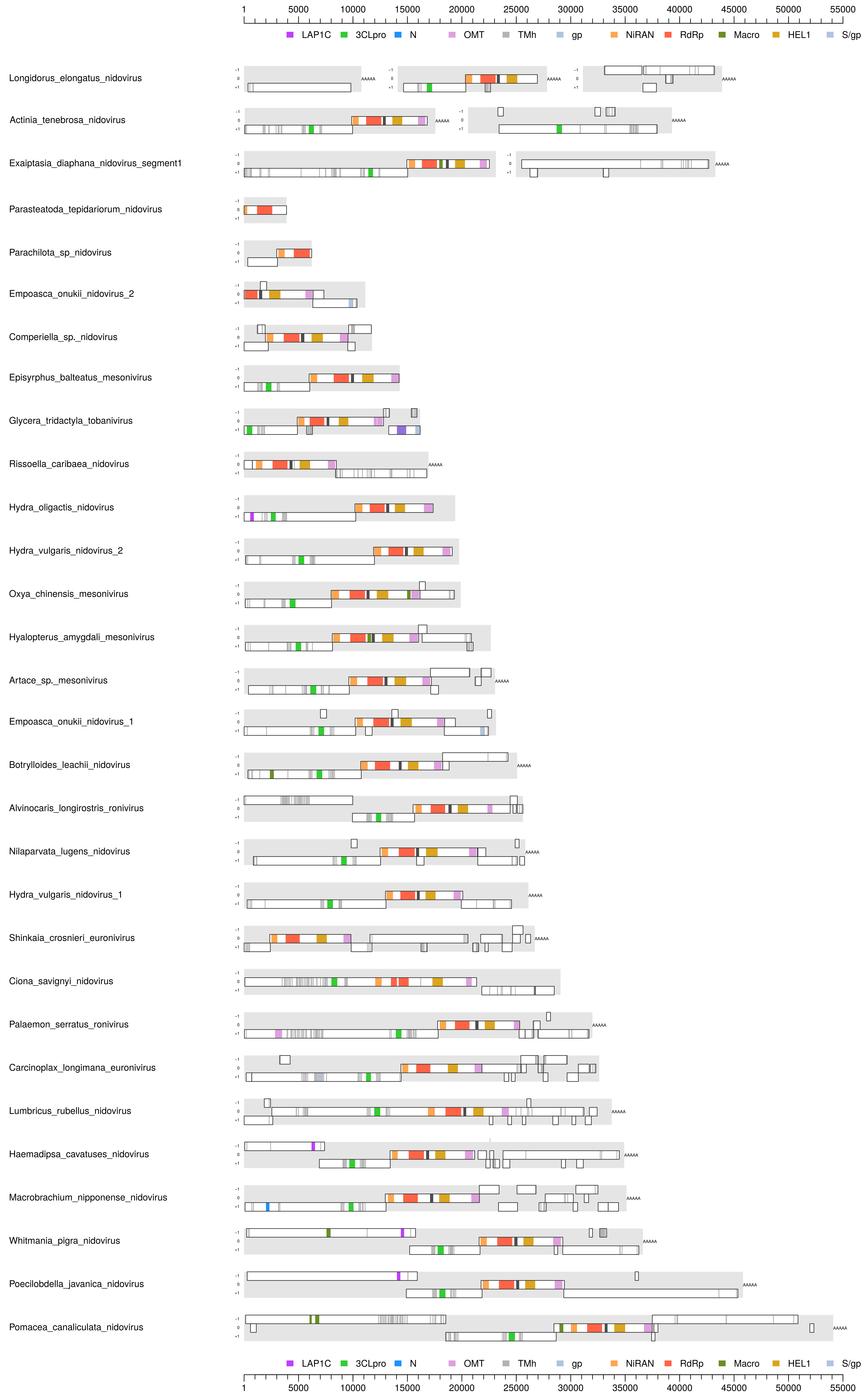

LAP1C 3CLpro N OMT TMh gp NiRAN RdRp Macro HEL1 S/gp
1 5000 10000 15000 20000 25000 30000 35000 40000 45000 50000 55000

### Figure S5

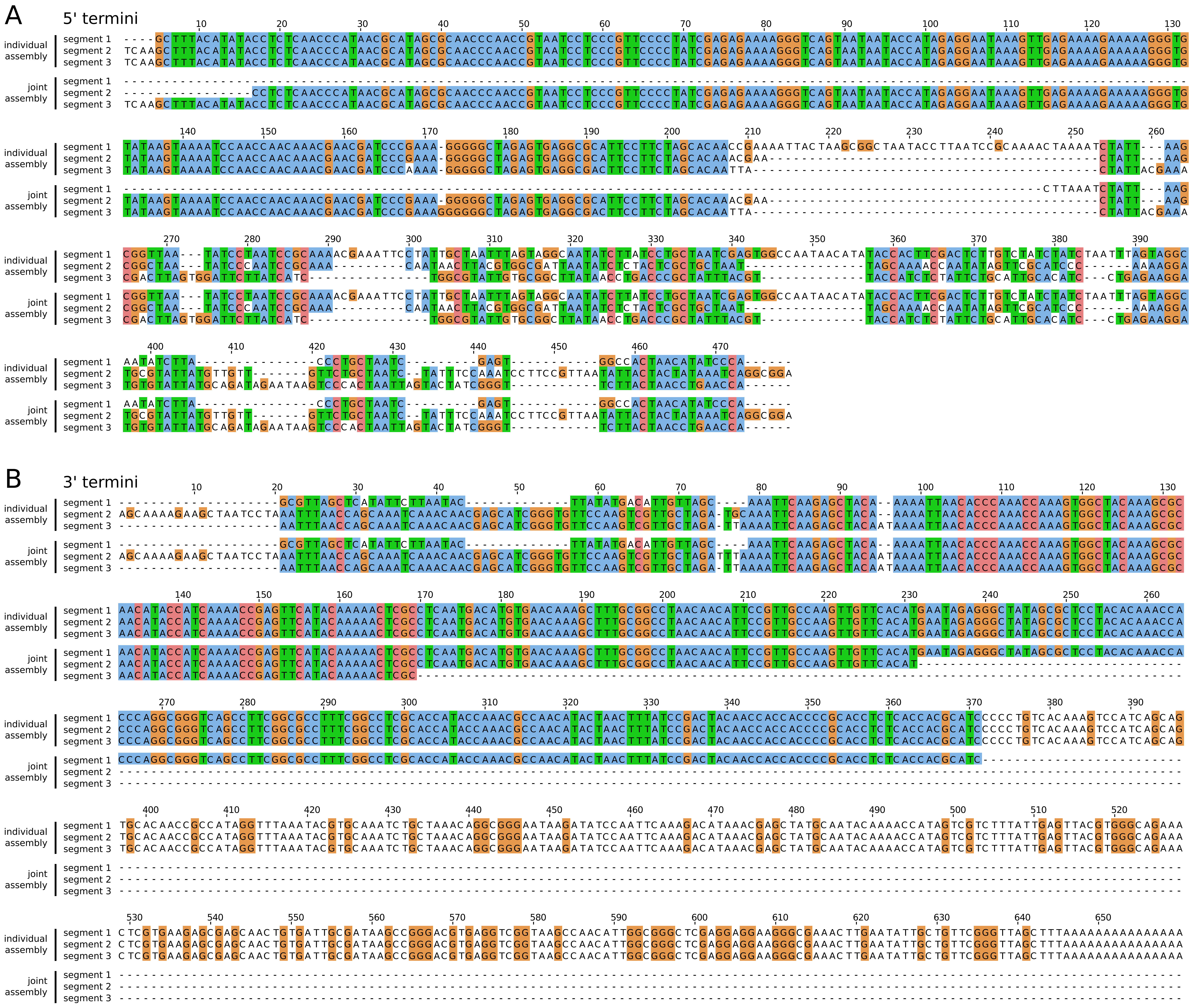

### Figure S6

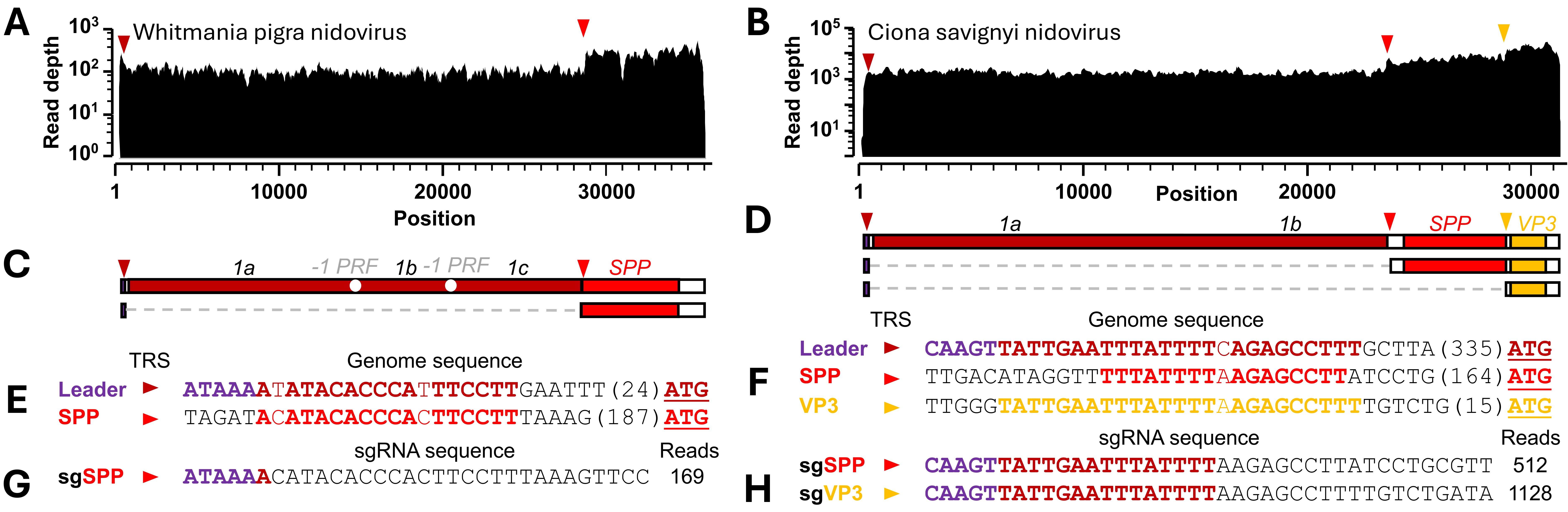
